# Applying Citizen Science to Gene, Drug, Disease Relationship Extraction from Biomedical Abstracts

**DOI:** 10.1101/564187

**Authors:** Ginger Tsueng, Max Nanis, Jennifer T. Fouquier, Michael Mayers, Benjamin M. Good, Andrew I Su

## Abstract

Biomedical literature is growing at a rate that outpaces our ability to harness the knowledge contained therein. In order to mine valuable inferences from the large volume of literature, many researchers have turned to information extraction algorithms to harvest information in biomedical texts. Information extraction is usually accomplished via a combination of manual expert curation and computational methods. Advances in computational methods usually depends on the generation of gold standards by a limited number of expert curators. This process can be time consuming and represents an area of biomedical research that is ripe for exploration with citizen science. Citizen scientists have been previously found to be willing and capable of performing named entity recognition of disease mentions in biomedical abstracts, but it was uncertain whether or not the same could be said of relationship extraction. Relationship extraction requires training on identifying named entities as well as a deeper understanding of how different entity types can relate to one another. Here, we used the web-based application Mark2Cure (https://mark2cure.org) to demonstrate that citizen scientists can perform relationship extraction and confirm the importance of accurate named entity recognition on this task. We also discuss opportunities for future improvement of this system, as well as the potential synergies between citizen science, manual biocuration, and natural language processing.

## Introduction

Biomedical literature is growing at rate of over a million new articles per year in PubMed (“PubMed Help” 2017) and represents a treasure trove of knowledge that forms the foundation for the design of future experiments. Making that knowledge more accessible and computable could save researchers time, effort, and resources (Yang et al., 2016) (Zhu et al., 2013). Researchers have effectively mined slices of the biomedical literature to identify potential treatments for Raynaud’s syndrome (Swanson 1986), drug candidates for Alzheimer’s disease (Li, Zhu and Chen, 2009), and potential mechanisms of ovarian oncogenesis (Urzúa et al., 2010). Given the large potential to make valuable inferences and the large volume of literature, many researchers have turned to information extraction algorithms to harvest information in biomedical texts and improve the value of existing data resources (Murray-Rust, 2017) (Pletscher-Frankild et al., 2015). Information extraction as a process can be divided into a few sub tasks: 1. Named Entity Recognition (NER), 2. Entity Linking (EL), and 3. Relationship Extraction (RE).

NER entails identifying specific types of entities within biomedical text (e.g., NGLY1 is a gene). Once identified, the NER term must be linked to an appropriate entry in a known database in order to provide semantic context (e.g., NGLY1 can be linked to gene 55768 in the NCBI Gene database). The process of linking NER annotations to known databases in order to provide context and generate semantic annotations is known as EL or normalization (Morgan et al., 2008) (Jovanović and Bagheri, 2017). After NER and EL, the relationships between the semantic annotations are extracted --e.g., [NGLY1] (gene) [mutations cause](relationship) [congenital disorder of deglycosylation](disease).

Algorithms for NER and EL have steadily improved thanks to the BioCreative challenges and availability of gold standard corpora (Wei, Kao and Lu, 2015). Tools such as EXTRACT (Pafilis et al., 2016) and those included in the PubTator suite (Wei, Kao and Lu, 2013) are sufficient for use in facilitating manual biocuration efforts. Furthermore, NER and semantic annotation algorithms have expanded beyond the concept types originally explored by the BioCreative challenges and now include post-translational modifications (Sun, Wang and Li, 2017), Gene Ontology terms (Ruch, 2016), metadata (Panahiazar, Dumontier and Gevaert, 2017), adverse effects (Cañada et al., 2017), and more (Tseytlin et al., 2016). Semantic annotation algorithms such as SemRep have been used to generate SemMedDB, a PubMed-scale repository of subject-predicate-object triples (Kilicoglu et. al., 2012).

Because of its dependency on NER, EL, and the limited availability of training corpora and ontologies, the automated approaches for RE have yet to reach the performance levels of NER and EL. To overcome those limitations, researchers have focused on improving NER and EL methods (Xing et al., 2018), different learning and modeling approaches (Peng et al., 2018), and expanding training data sets via mining of knowledge bases (Zhou et al.) or crowdsourcing via paid microtask platforms (Lossio-Ventura et al.)(Li et al., 2016). Crowdsourcing through paid microtask platforms to expand the training data sets has proven to have great potential, but questions regarding scalability prompted us to investigate citizen science as a potential avenue for crowdsourcing RE.

Citizen science is a form of crowdsourcing in which non-professional scientists voluntarily engage in different degrees of data collection, analysis, and/or dissemination of a scientific project (Haklay, 2013). The scalability of citizen science has enabled researchers to collect, process, and analyze unprecedented volumes of data leading to advances in conservation and environmental science (Schmiedel, 2016)(McKinley et al., 2017), astronomy (Banfield et al., 2016)(Kuchner et al., 2016)(Straub, 2016), biomedical research (Candido dos Reis et al., 2015) (Luengo-Oroz, Arranz and Frean, 2012)(Kim et al., 2014), and more (Williams et al., 2014)(Palermo et al., 2017). Crowdsourcing and citizen science has previously been applied towards NER of disease mentions via a platform called Mark2Cure (M2C), and found that in aggregate, annotations submitted by trained citizen scientists were on par with expert annotators (Good et al., 2015) (Tsueng et al., 2016). Based on this finding, citizen scientists may serve as an additional check for annotations generated by computer algorithms, and address quality issues introduced by NER and EL tools. The problem of insufficient gold standard corpora for RE tasks can also be addressed by crowdsourcing (Aroyo and Welty, 2013). In aggregate, non-experts recruited via a microtask platform could perform relationship annotation on par with that of a single expert annotator (Dumitrache, Aroyo and Welty, 2015).

In this paper we describe the application of citizen science towards RE from biomedical text. Specifically, we 1. Provide a brief overview of the RE module within the Mark2Cure platform, 2. Evaluate the ability of citizen scientists to perform RE taking into consideration the limitations and ambiguities inherent in the system, and 3. Compare the citizen science generated data with the automated results from SemMedDB to understand how the two may complement and enhance each other.

## Methods

### Mark2Cure Relationship App Design

The NER module within the Mark2Cure platform has previously been described (Tsueng et al., 2016). In addition to the NER module and shared dashboard, there is a RE App which has a separate training module, task list, task interface, and feedback screen. Mark2Cure is an open-source project, and code is available at https://github.com/SuLab/mark2cure.

For the RE App, training consists of a series of interactive modules developed after several iterations of testing and feedback from the Mark2Cure community. The first module introduces the user to the task interface. The second module introduces the user to the 2 fundamental rules of the task: 1. select relationship based only on what is in the abstract (no prior knowledge), 2. select most granular relationship without guessing. The third module has three submodules introducing the user to the different kinds of relationships that they will need to classify (gene-disease relationships, gene-treatment relationships, and disease-treatment relationships). Because there are no gold standards for this task, users are provided with visual feedback on how their selection aligned with that of the all the other users who have done the same task. Each task needs to be evaluated by multiple users in order to be considered complete.

The Mark2Cure relationship app currently pulls concept annotations using the PubTator suite (Wei, Kao and Lu, 2013). For each abstract, every combination of heterogeneous concept pairs is calculated and designated as a task. Concept pairs within the same concept type are not included because the relationship between concepts of the same concept type tend to be hypernymic relationships (e.g.-“is a”) which algorithms are good at identifying (Rindflesch & Fiszman, 2003). Users are presented with a concept pair and the concepts are highlighted in the abstract to provide context. Based on the abstract text, the users are asked to identify the type of relationship (or lack of relationship) between those two concepts. Users also have the ability to tag either of the concepts as incorrectly identified/inappropriately annotated.

Mark2Cure is an ongoing project with active data collection and user submissions. The data analyzed for this study was collected between 2016.05.26 and 2017.11.22. This data set consists of 4047 concept pairs pulled from 1058 abstracts annotated by 147 contributors; of which 1009 concept pairs from 234 abstracts were marked by at least 15 different contributors. The data set used in this analysis can be found at https://github.com/gtsueng/M2C_rel_nb/data.

### Analytical methods

#### Data and code availability

All the code and data used in the following sections can be found at https://github.com/gtsueng/M2C_rel_nb. Figures were generated as coded in the jupyter notebooks using the bokeh library (if interactive).

#### Generation of a M2C-specific pseudo-gold standard for quality control (QC)

There were 1009 completed concept pair relationship (i.e.-task) annotations at the time of analysis, and a 10% sample (~100 concept pairs) was desired for quality control. 120 PubMed ID (PMID) PMID-specific concept pairs were randomly selected and manually inspected to determine the expected response based on the rules and available options.

#### Contribution distribution, accuracy, and aggregation threshold determination

The contribution distribution was limited to just the set of completed task annotations. For this set, the number of RE task annotations that each user contributed was determined, sorted, and plotted. For individual accuracy estimation, the set of each user’s task annotations was compared with the QC annotation set. The accuracy was estimated based on the intersecting task annotations of the two sets and the median of all user accuracy estimations was calculated. For aggregate accuracy estimation, the user annotations for the concept pairs that were QC’d were pulled into a dataframe for further analysis. User agreement thresholds (K) were set at 1 (no agreement) to 15 (max agreement). For each concept pair, at each value of K, K users that annotated that concept pair were randomly selected and the majority response for that concept pair was identified. If the results were tied, one of the tied responses were selected at random. For each value of K, the random selection and majority determination was performed 10 times (i.e.-10 iterations). To stabilize the results, the entire process was repeated 5 times for each concept pair (i.e.-5 repetitions). The accuracy of the responses in the QC’d set were calculated per each level of k, repetition, and iteration. The median accuracy for each level of K was calculated for each repetition and then averaged over the number of repetitions.

#### Identification of missing relationship types and verification of non-relationships

To identify missing relationship types, PMID-specific concept pairs annotated as ‘has relationship’ or ‘other relationship’ were aggregated to obtain the total number of users that marked each concept pair as having a non-specific relationship. At each voter threshold (K), up to 25 PMID-specific concept pairs were randomly selected for qualitative review. The concept pairs and the respective user counts were exported, randomly assigned a number and randomly sorted by that number. The user counts were then masked to prevent biasing and each PMID-specific concept pair was reviewed in-house. The same process was applied to PMID-specific concept pairs marked as having ‘no relationship’ or ‘cannot be determined’.

#### Evaluating the effect of concept distance on accuracy

This analysis was run on only the relationship annotations data for only completed concept pairs, for which every concept has an associated identifier. The abstracts were tokenized at the sentence level using the NLP Tool Kit (NLTK) (Bird, Loper, and Klein, 2009) to obtain an average per-sentence character count which can be used to estimate the concept distance at the sentence level. Only concepts with known identifiers were analyzed as it would be more difficult to determine the positional location of a term in an abstract when the identifier is missing. Since discarding the concept annotation should not be affected by concept distance, relation annotations for concepts considered correctly annotated (i.e-’not broken’) were treated separately in this analysis.

#### Comparison with PMID-specific SemMedDB relationships

The PMIDs for the completed concept pairs were used to pull SemMedDB annotations specific to those PMIDs. SemMedDB concept annotations are linked to Concept-Unique Identifiers (CUIs) from the Unified Medical Language System (UMLS). The UMLS integrates many biomedical vocabularies and standards (Bodenreider, 2004). Since UMLS supports many more concept types than Mark2Cure, only UMLS concept types that mapped to Mark2Cure concept types were included in the analysis. Additionally, the RE module in Mark2Cure does not yet appear to be connected to the NER module; hence, the Mark2Cure relationship concepts are purely based on PubTator. Thus, the UMLS concept types were mapped to Mark2Cure concept types based on mappings used in in the generation of the corpora used for the gene, disease, and drug NER algorithms in PubTator with a few additional mappings to suit the expansion of the PubTator concept types (Mark2Cure Relation Extraction concept types) to the Mark2Cure NER concept types. The identifiers for relationships considered complete in Mark2Cure were converted to UMLS CUIs and used to filter an export of SemMedDB annotations for only relationships involving those CUIs. The semantic types of the SemMedDB subject and objects were mapped to Mark2Cure concept types for comparing relationships within the same entity type vs different entity types. The types of concept pairs were compared between SemMedDB and Mark2Cure after concept pairs in which the majority of users marked a concept as incorrectly identified (i.e.-‘broken’) or unrelated were removed from the remaining set of PMIDs.

## Results

### Contribution distribution, accuracy, and aggregation threshold determination

We used Mark2Cure to annotate the relationships between 1009 concept pairs from biomedical abstracts, and each pair was annotated by at least 15 Mark2Cure volunteers. In total, we collected and analyzed 15,739 annotations from 147 volunteer contributors. As with other crowdsourcing systems, the observed distribution of contributions was unequal. The Gini coefficient of the distribution of contributions plot (Figure 1A) was 0.73 which was comparable to what was observed in the disease mentions NER pilot study (gini = 0.716)(Tsueng et al., 2016).

**Figure 1A -.**
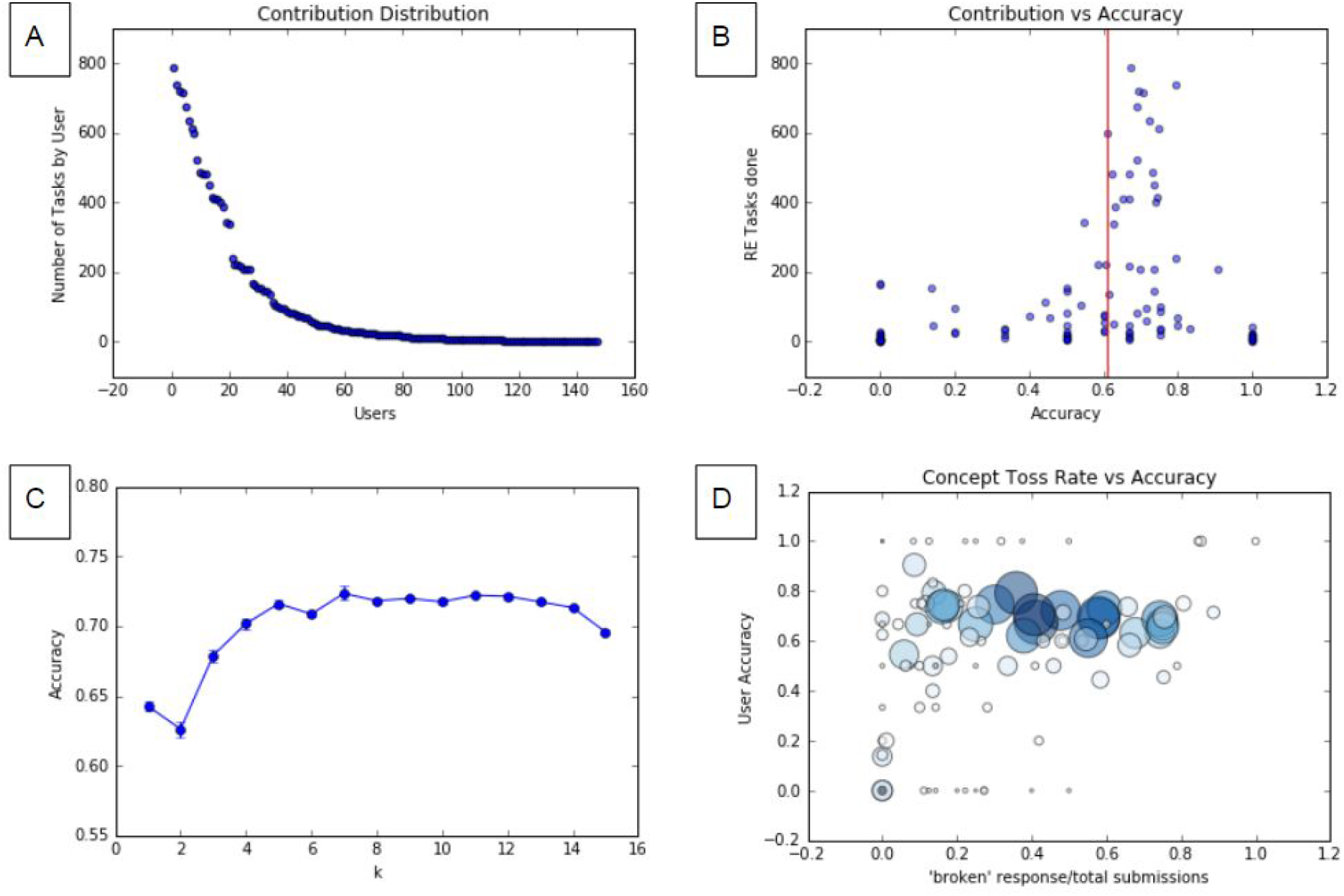
Contribution distribution of the RE task. B. Each user’s estimated accuracy (x) vs the number of tasks that user completed (y). The red line illustrates the median accuracy. C - Median accuracy with respect to the majority response of an aggregate of K users. The concept toss rate (i.e.-concepts marked as broken) vs accuracy of individual users (D). The size of the circles in D represent the number of total tasks that individual user contributed, while the color represents the number of that user’s annotations which could be found in the QC set and therefore used for accuracy estimations.

To assess the accuracy of these relationship annotation results, we compared them to a manually-curated subset of the full data set. Based on this quality control (QC) set, the median accuracy per user across all of their annotations was 0.61 (Figure 1B).

To assess how increasing the number of contributors affected the quality of the aggregate annotations, we simulated smaller numbers of annotators per document by randomly sampling from the QC set. When aggregating user responses and selecting the majority response, the accuracy increases along with the number of users K up until about 6 users. Beyond 6 users, the median accuracy does not further increase, suggesting that each relationship task should be annotated by a maximum of six users in order to maximize accuracy while minimizing redundancy of work done by the community (Figure 1C).

A small subset of users were estimated to have very low accuracy despite contributing over ten task annotations--warranting the need to verify that there weren’t unaccounted for issues with the data. Upon manual inspection of two of these outliers, we observed that these users did not mark any concepts as inappropriately annotated. To investigate the effect of a user’s reluctance to correct incorrectly identified concepts, we calculated each user’s ‘concept toss rate’ (i.e.-number of annotations they marked as incorrectly identified relative to number of annotations they submitted) and plotted their estimated accuracy relative to their toss rate (Figure 1D). A subset of users within the lowest range of accuracy also had a low average rate of tossing out concepts highlighting the effects of NER quality on Relationship Annotation. Inquiries sent from a few users suggested that at least some of the error could be attributed to a lack of guidance on how to prioritize when dealing with multiple true states. For example, an incorrectly identified concept can be correctly marked as inappropriately annotated, but also correctly marked as having ‘no relationship’ since the incorrectly identified concept is not under discussion in the abstract. Further clarification on how to prioritize multiple true responses could help to improve consistency and performance of RE across the Mark2Cure community.

### Identification of missing relationship types

The relationship annotation options in Mark2Cure were based on higher level relation properties from an ontology in development in WebProtoge (“`WebProtégé”, n.d`.). With limited relationship options available in Mark2Cure, qualitative analysis of concept pairs annotated as ‘has relationship’ or ‘other relationship’ can provide insight into relationships missing from the currently available options in Mark2Cure. To identify missing relationship types, PMID-specific concept pairs annotated as ‘has relationship’ or ‘other relationship’ were aggregated to obtain the total number of users that marked each concept pair as having a non-specific relationship. We sampled up to 25 PMID-specific concept pairs at each voter threshold (K), randomized the order of the samples, masked the number of users that marked it as having a non-specific relationship, and then manually inspected the relationship. If the relationship between the two concepts was an available option within the Mark2Cure system, it was binned as ‘has available specific relationship’. If either of the concepts were inappropriately annotated, it was binned as ‘concept broken’. Common relationships not available in the Mark2Cure system were binned together and new categories were created whenever the a relationship did not fit in with previous categories.

As seen in Figure 2, there is a decreasing number of per-PMID concept pairs that were marked as having a better response option within Mark2Cure (pink/red squares) as the number of users that agreed with that assessment increased. Per-PMID concept pairs that were marked as ‘has relationship’ by a high number of users tended to genuinely have a relationship not captured by the system. However, setting the threshold too high increases the risk of missing interesting, non-obvious relationships that are otherwise not captured by Mark2Cure.

**Figure 2 -.**
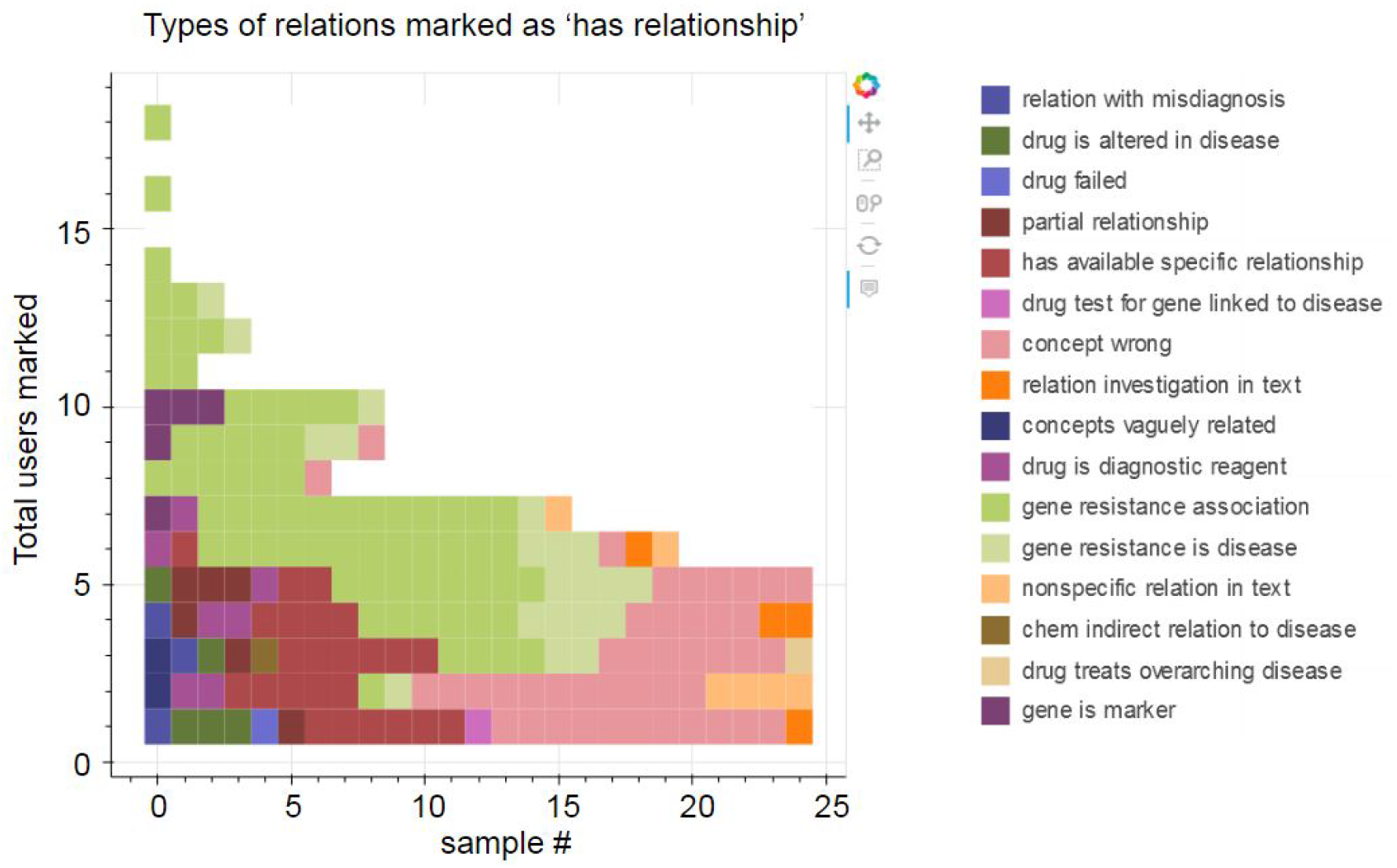
Qualitative assessment of a sample of relationships marked as generic relationship/other relationship by total user counts. In pink are ‘has relation’ annotations for a concept pair in which one of the concepts appear to be incorrect (i.e.-broken). In red are ‘has relation’ annotations for concept pairs which could be described with a more specific option. In green are concept pairs with a genuine relationship, not available as a selection option. All other colors indicate other ways in which the two concepts are discussed together (e.g.-purple: gene is a marker used to study samples from patient with disease, etc). For the fully interactive Figure 2, visit (https://github.com/gtsueng/M2C_rel_nb/tree/master/results/has_relation_broad_categories.html).

Interesting relationships that were missing from Mark2Cure’s selection options included: Resistance/Insensitivity to the gene was associated with the disease; the disease conferred resistance to the drug; the drug was altered in the disease; the gene is a marker for inspecting samples involving the disease; the drug is used in the diagnosis of the disease; and a mutation in the gene causes a disease that was misdiagnosed as the disease. Some users also marked ‘has relationship’ for concept pairs which had an explicitly inconsistent relationship in a few case studies or when the text explicitly mentioned the investigation of a relationship between two concepts without actually revealing the relationship. Since users in aggregate appear to be correctly identifying missing relationships as ‘has relationship’, we investigated the annotations marked as ‘no relationship’ to understand any rules/guidelines in the system in need of further clarification.

### Verification of unrelated concepts and identification of rules in need of improvement

We applied the method used for Figure 2 to PMID-specific concept pairs that users marked as having ‘no relationship’. Few PMID-specific concept pairs (each RE task) were marked by at least six users as having ‘no relationship’; hence, sampling the RE tasks for qualitative analysis was only necessary for RE tasks with five or less users agreeing on the ‘no relationship’ response. At K=6, the number of different RE tasks that were marked as having ‘no relationship’ drops to only seven; and drops again to less than half of that at K=7. At K=8 or above, we were unable to find RE tasks marked by exactly K users for each level of K above 8, severely limiting the sample size (Figure 3).

**Figure 3 -.**
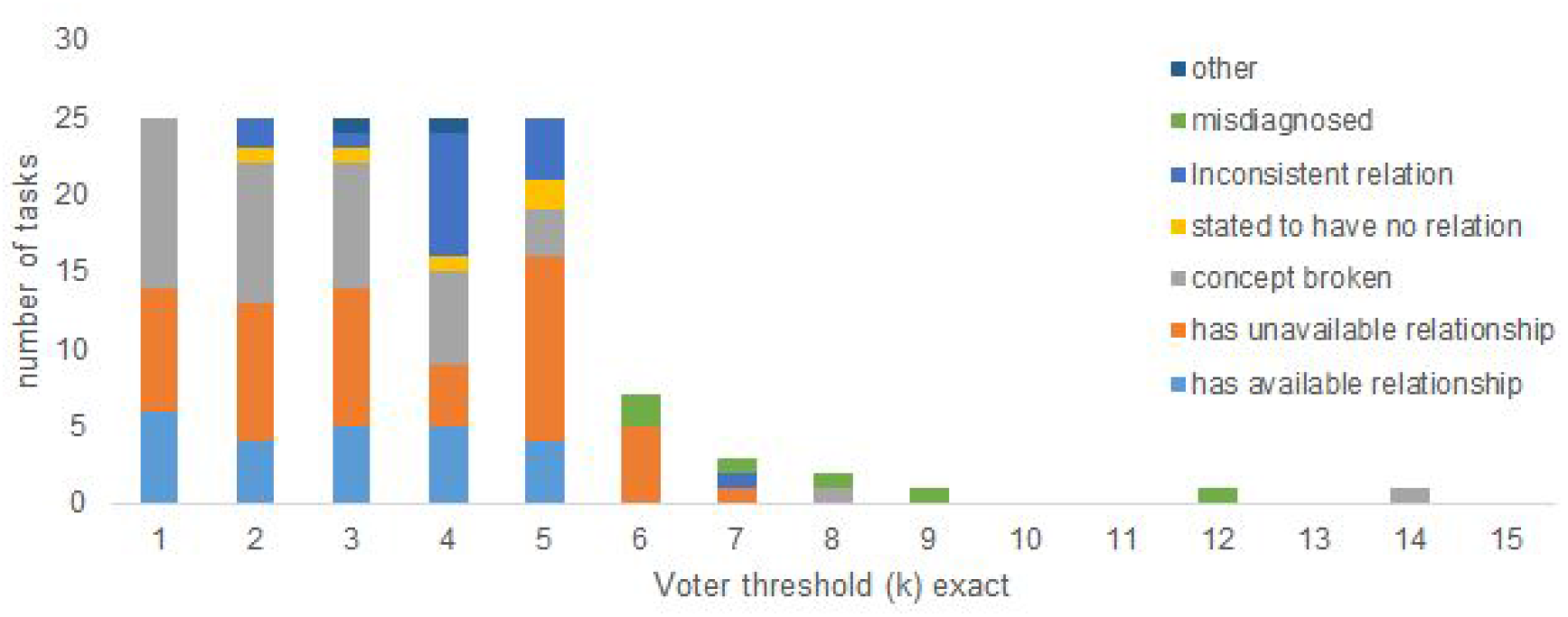
Types of relationships marked as having ‘no relationships’. Light blue indicates that an appropriate relationship was available in the system even though the majority ruled there to be ‘no relationship’. In orange are concept pairs determined to have a relationship that is currently not available in the system. Grey indicates concept pairs in which one of the two are considered incorrectly annotated. Blue indicates that the text suggests an inconsistent or partial relationship between the concept pairs; hence, the concepts can simultaneously have ‘no relationship’ and a ‘specific relationship’. Green indicates that the relationship is between a gene or drug and a disease which was misdiagnosed in lieu of the actual disease. Other types of relationships are in dark blue.

For the qualitatively inspected tasks--at each value of K that was below six, a better relationship (generic or specific) existed 20% of the time. At each value of K of six or above, a better (generic or specific relationship) was not available. As K increases from 1 to 5, the number of tasks in which a concept should have been marked as inappropriate decreased from 11 to 3. At K>=6, the number of tasks in which a concept should have been marked as inappropriate was mostly 0. RE tasks for which a specific relationship exists (but the option is not available) averaged to make up about a third of the annotations marked as having no relationship when K was less than 6. Instances where a drug failed to treat a disease and therefore can truly be counted as having ‘no relationship’ was not observed when at K<5 due to the limited sample availability. Misdiagnoses accounted for some of the instances of ‘no relationship’ for RE tasks with K>=6. See Supplemental Figure 1 for the breakdown and variety of ‘no relationship’. Based on these results, clarification and improved guidance is needed for the treatment of abstracts that cover case studies or clinical trials (in which the relationship may be inconsistent/partial) and for instances where a relationship was suspected, but later found to be untrue such as in a misdiagnosis or failed drug trial.

### Effects of concept distance on relationship identification

Many semantic and NER text mining algorithms perform optimally when used in conjunction with text tokenized at the sentence level (Muzaffa, Azam, and Qamar, 2015)(Lou et al., 2017) (Zhu et al., 2017). Limiting the Mark2Cure RE task to concepts that share the same sentence could reduce the amount of text contributors would need to read and reduce the amount of text displayed in mobile devices. However, such limitations would also result in losing the option to identify relationships between concepts in the text that do not appear in the same sentence. To evaluate the advantages and disadvantages of restricting the task to the sentence-based concept pairs, we looked for relationship annotations of concepts that were not in the same sentence and we inspected the effects of the distance between concept pairs in a task on accuracy.

As seen in Figure 4, most concept pairs were less than a sentence apart. Multiple synonymous mentions of concepts increase the likelihood of concept pairs to be located within shorter distances than farther ones, nonetheless there were still plenty of relationship identified between concept pairs that were estimated to be two or more sentences apart. In cases where the concept distance was not expected to affect the relationship (i.e.-one of the concepts is considered inappropriately annotated), there were still more concept pairs located closely together in space than farther apart. Estimated accuracy appeared to be more affected by voting threshold (K) than by the estimated minimum distance between the concept pairs. This was particularly visible in the difference observed at K=15 between concept pairs with a relationship and concept pairs marked as broken (Supplemental Figure 2). For concept pairs considered broken, the accuracy at K=15 decreases due to the inclusion of annotations from users uncomfortable with discarding concepts as broken in all runs. The concept pairs were further subdivided between those marked as ‘unrelated’ in the QC set to determine if unrelated concept pairs were more likely to be located farther apart. This was not found to be the case (Supplemental Figure 2). Note that not every annotation from Pubtator was linked to an identifier. Annotations lacking identifiers may or may not be synonymous with other annotations lacking identifiers within an abstract, making it difficult to calculate the minimum distance within the text between two concepts if there are multiple annotations lacking identifiers.

**Figure 4 -.**
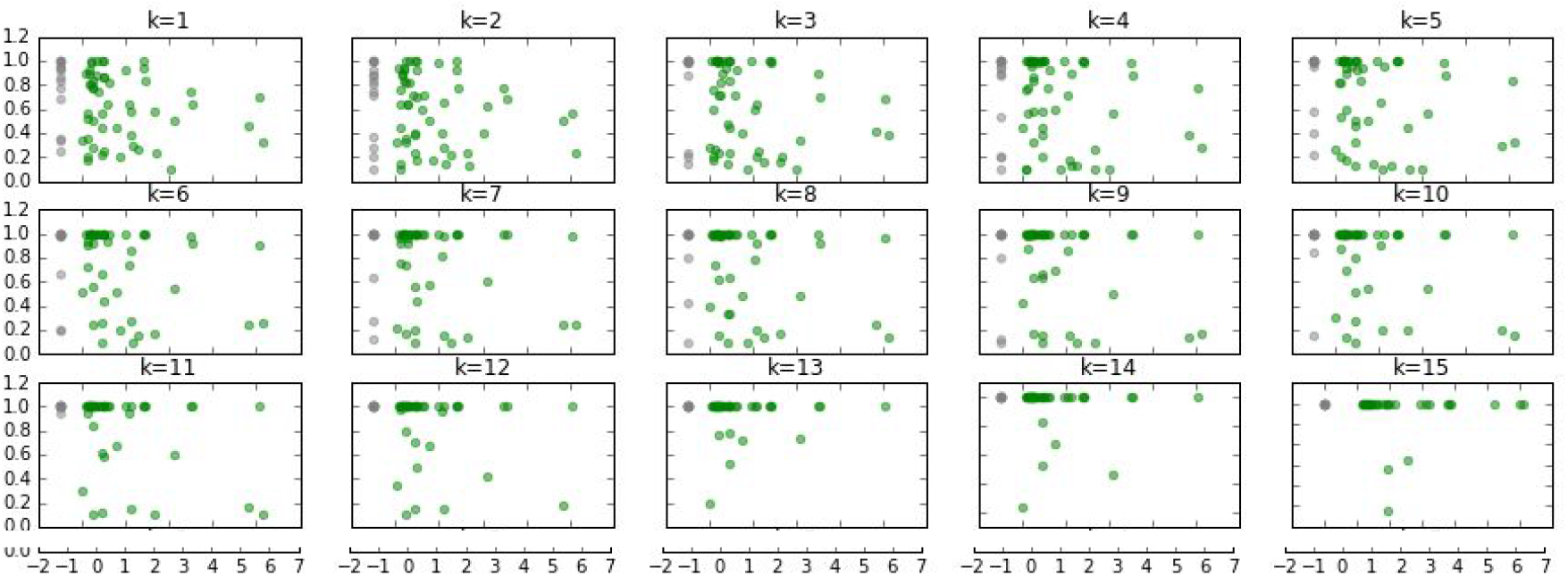
The minimum estimated number of sentences between two concepts (x-axis) vs the average accuracy of the majority response (y-axis) at different voter thresholds (K) for concept pairs which were not considered inappropriate (green). In grey are concepts for which no identifier was available and the minimum distance was not calculated.

### Comparison with PMID-specific SemMedDB relationships

Although concept distance is one factor which distinguishes Mark2Cure from automated methods that tokenize at the sentence level, we wanted to investigate how the relationships annotated in Mark2Cure compared with those annotated via automated methods. We pulled subject-predicate-object triples from SemMedDB and restricted the SemMedDB data to just the abstracts in common with the Mark2Cure data set. Some differences in the relations between SemMedDB and Mark2Cure were immediately visible. SemMedDB tokenizes at the sentence level and mines the relationships from these sentences regardless of concept type resulting in a very different set of relationship annotations as compared to Mark2Cure. In contrast, Mark2Cure users are only presented with concept pairs which are different in type, no matter where they may appear in the abstract. For example, users may be asked about the relationship between a gene term and a disease term, but never about the relationship between a disease term and a different disease term. Additionally, SemMedDB has many more entity types than Mark2Cure, and in order to do a more detailed comparison, we restricted our comparison of the two to the entities that were found in common.

As seen in Figure 5, the majority of relationships in SemMedDB for the abstracts that have at least one subject entity and one object entity in common with Mark2Cure are relationships within the same type of concepts. Only four abstracts contained concept pairs that appeared to have a relationship in both SemMedDB and Mark2Cure. Most of the relationships in Mark2Cure appear to be gene-disease relationships, but this is due to the removal of concept pairs which were marked as inappropriately annotated. Given the differences in relationship types available in SemMedDB and Mark2Cure, we wanted to explore the potential complementarity between the two. For example, gene-disease relationships which are likely to be underrepresented in SemMedDB due to the sentence-level tokenization could appear with greater frequency in Mark2Cure. Similarly, broad/non-specific relationships identified in Mark2Cure could be explained by multi-node (indirect) relationships or other relationships available in SemMedDB but not in Mark2Cure.

**Figure 5A -.**
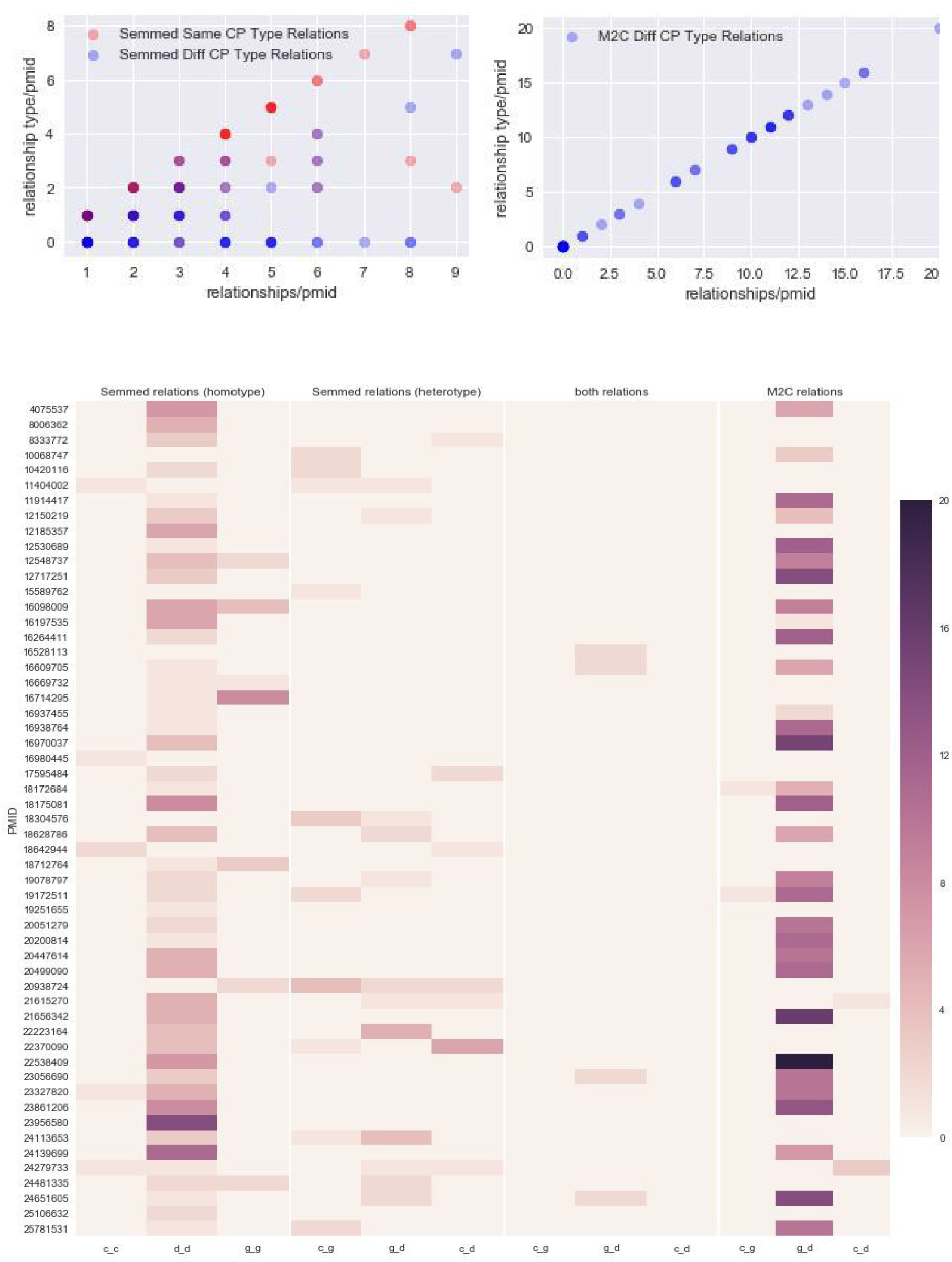
Distribution of types of concept pairs per abstract vs the total number of relationships per abstract. Since Mark2Cure only allows relationships between different types of concepts, the number of relationships for different types of concepts per abstract will be the same as the number of relationships per abstract regardless of type (right). In SemMedDB, the number of relationships of similar concept type per abstract (red) is higher than the number of relationships of different concept type per abstract (blue) resulting in the distribution on the left. 5B-Heatmap illustrating the number of relationships identified in each abstract via SemMedDB, both SemMedDB and Mark2Cure, and just Mark2Cure. In the case of Mark2Cure only true relationships (broad or specific) were counted (inappropriately annotated concepts or ‘unrelated’ responses were not included in the count).

To explore the potential complementarity of the data from the two systems, the quality-checked concept pairs were aggregated in terms of the majority response given per PMID in Mark2Cure. The concept pairs were also checked for a direct relationship and for co-occurence of the concepts in the pair (potential indirect relationships) in SemMedDB after filtering for only relationships within the quality-checked PMIDs. As seen in Figure 6, concept pairs with specific relationships appear in many more PMIDs in Mark2Cure than in SemMedDB. This difference is largely attributable to NER differences between PubTator and SemMedDB.

**Figure 6 -.**
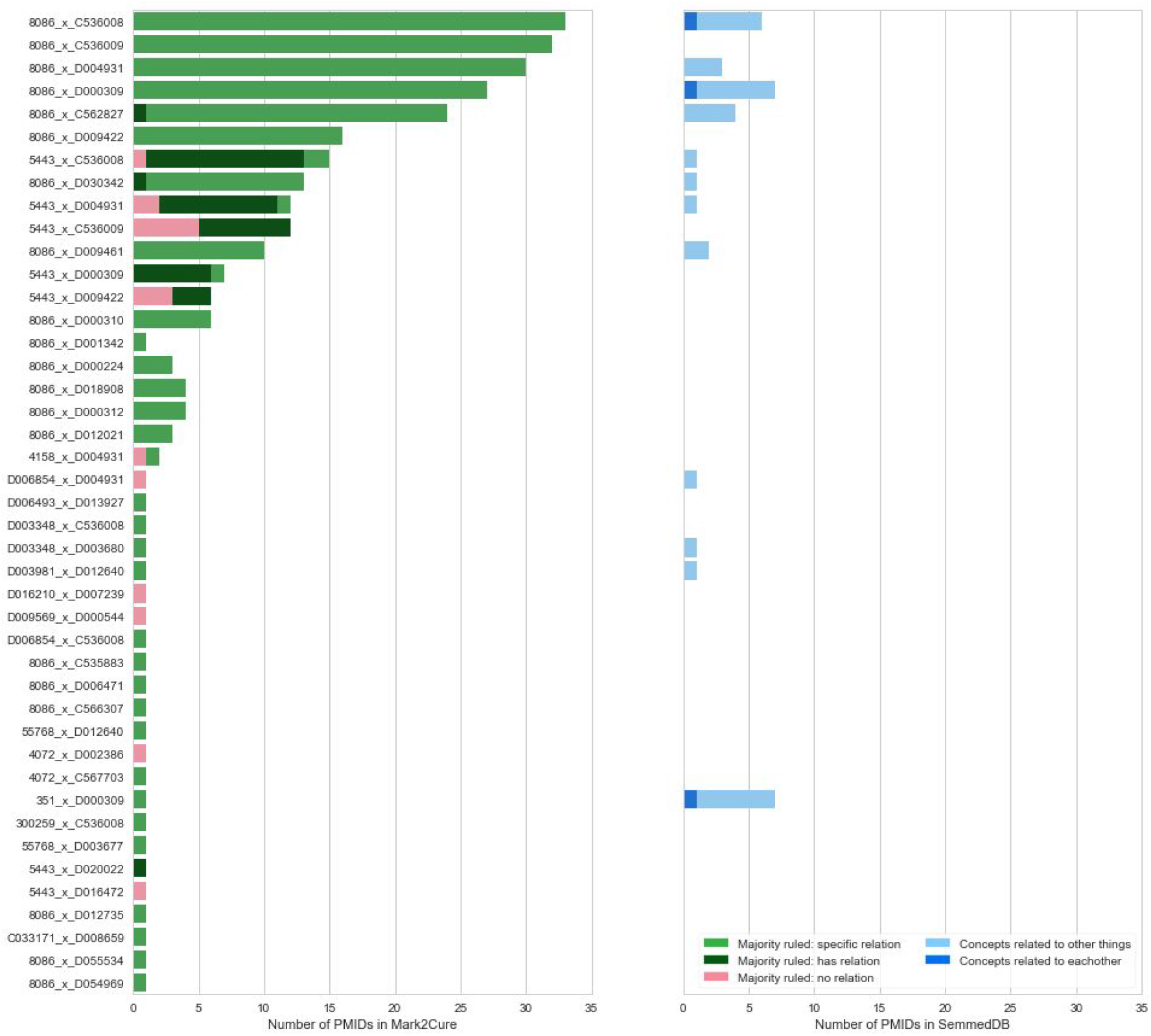
Within the set of concept pairs in the QC set, the number of PMIDs in which the concept pairs appear within the set of abstracts completed by Mark2Curators is on the left along with the majority response for that concept pair across different abstracts. In green are the number of abstracts in which the majority of users ruled that the concepts had a specific relationship, dark green is the number of abstracts in which the majority of users ruled that the concepts had a nonspecific relationship, and in pink are the number of abstracts which the majority of users ruled that the concepts were unrelated. On the right are the same concept pairs pulled from SemMedDB restricted to the same set of abstracts as they appear in relationship with other concepts (light blue) or in a direct relationship with one another (blue)

For example, the concept pair 8086_x_D000309’ (AAAS gene x Adrenal Insufficiency) was observed in 28 abstracts (when annotated by PubTator), but only 6 of those abstracts were both entities part of relationships in SemMedDB. In some cases these were due to differences in the way text was annotated by PubTator vs SemMedDB (e.g.-PubTator marking mentions of ‘adrenal insufficiency’ as mentions of ‘Triple A syndrome’ or ‘Allgrove syndrome’). In other cases (like ‘8086_x_D009461’: AAAS gene x neurologic dysfunction), mentions observed in PubTator were missed altogether in SemMedDB. If each of the concepts in a concept pair were found in an abstract in SemMedDB, the relationship between the two concepts was not annotated unless the two concepts were in the same sentence as seen in the case of D003981_x_D012640’ (Diazoxide x Seizures). This concept pair was observed in PMID: 11916319 by both PubTator/Mark2Cure and SemMedDB; however, because the two concepts were found in different sentences, the relationship between the two concepts is not annotated in SemMedDB.

The frequency of relationships in concept pairs arising from Mark2Cure may be useful for selecting important relationships in SemMedDB. Data from Mark2Cure can be used to identify relationships missed from SemMedDB due to sentence-level tokenization. Furthermore, SemMedDB may be useful for clarifying relationships between concept pairs that users consistently marked as having an unspecified relationship if the concepts in Mark2Cure can successfully be mapped to those in SemMedDB.

## Discussion

We found that citizen scientists were willing and capable of performing relationship extraction (RE), and that the relationships they extracted from the full abstract was different than those obtained via automated methods like SemMedDB. These differences may be due to system-specific restrictions such as sentence-level tokenization in SemMedDB and inclusion of only heterogeneous concept pairs in Mark2Cure. Aggregate task performance by the citizen science community was affected by three primary issues: 1. NER quality issues, 2. training and platform issues, and 3. issues with the documents. Consistent with the literature on information extraction, the quality of the RE data was affected by the NER quality issues (Xing et al., 2018) (Li et al., 2016) and users who were uncomfortable or unwilling to discard concepts had lower performance results. In aggregate, the users generally selected the most appropriate response in spite of the NER issues and limited (and sometimes ambiguous) choice options in the RE task. In spite of multiple iterations in the development of the training and platform in an effort to ensure high quality data (Gabriele and Eva-Maria, 2016)(Kosmala et al., 2016), design issues such as the lack of guidance on prioritizing multiple true responses still affected the performance of the citizen scientists. We expect that addressing these issues will improve contributor performance.

By investigating concept pairs that had a high levels of agreement for different responses in different abstracts, we identified areas in need of refinement in terms of available relationship options (e.g. - missing relationships) and modeling (e.g.-inconsistent relationships, misdiagnoses); which led us to identify issues with the documents themselves. The majority of the documents from which the Mark2Cure community extracted relationships surround a more unique symptom of NGLY1-deficiency (alacrima). Hence the documents were more homogenous and narrow in scope, limiting the types of missing and non-relationships that could be identified. Furthermore, this document set had a number of case studies in which the relationship between a pair of concepts could be simultaneously true and false (e.g.-gene mutation was involved in a disease in 2 out of 3 patients). Based on our findings, we expect to be able to raise the community performance on the RE task by providing more guidance on reviewing the concepts (NER) and prioritizing responses in situations involving multiple ‘true’ responses and/or simultaneous ‘true and false’ responses. As with crowdsourcing relationship extraction (Aroyo and Welty, 2013), applying citizen science to relationship extraction is a potential avenue for generating new training data sets for improving automated relationship extraction tools.

Although preliminary, we demonstrate that citizen scientists can contribute different types of relationship annotations across three different types of concepts. In contrast, many relationship extraction efforts focus on a specific type of relationship extraction (Li et al., 2016) (Cañada et al., 2017)(Xing et al., 2018)(Collier et al., 2015). This difference makes it difficult to draw comparisons on contributor performance, but opens up interesting avenues for exploration in relationship extraction. In particular, it would be interesting to evaluate the results from this approach with those from non-specific (Mintz, Bills, Snow & Jurafsky, 2009) and medically-tailored (Wang & Fan, 2014) relationship extraction algorithms, or those from crowdsourced efforts involved in active (Liu et al., 2016) or semi-supervised (Angeli et al., 2014) learning.

## Conclusion

This experiment demonstrates the application of citizen science to relationship extraction and when taken together with the previous findings on disease NER (Tsueng et al., 2016), demonstrates that citizen science can be applied towards information extraction in biomedical abstracts. We discuss data quality issues and the training improvements that could be used to address them; and we compare the citizen science-generated data with automated, text-mined data to understand the differences between the two and how they may complement one another.

## Acknowledgments

This paper was made possible by citizen scientists. The importance of their contributions both within the Mark2Cure system and outside (especially their questions, feedback, suggestions) cannot be emphasized enough, and we are extremely grateful for the privilege of working with such wonderful contributors and collaborators. Contributors who consented to publishing their names can be found here: https://mark2cure.org/blog/relation-paper-contributors/. We would also like to thank the Mights, the Esteniks, the Jennisons, and other members of the NGLY1 rare disease community for their inspirational activities and words of support. This work was supported by the US National Institute of Health (U54GM114833 to A.I.S.). This work was also supported by the Scripps Translational Science Institute, an NIH-NCATS Clinical and Translational Science Award (CTSA; 5 UL1 RR025774).

## Appendix

**Supplemental Figure 1-.**
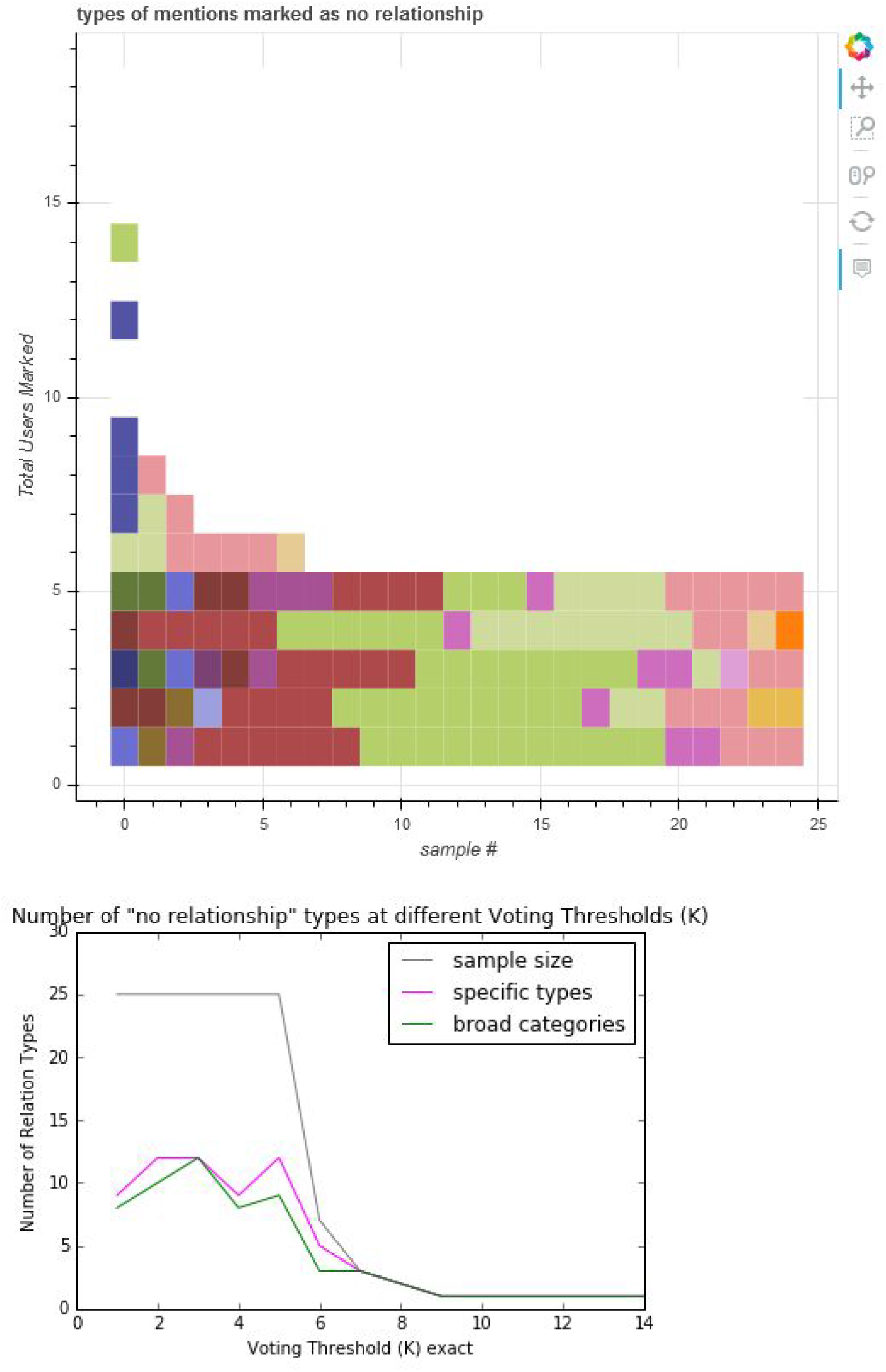
Types of concept mentions marked as ‘no relationship’ or ‘cannot be determined’. Red and pink: concept pairs that have some sort of relationship; green: concept pairs that are incorrect/should be thrown out. Light green: concept pairs which have a relationship only in x/y cases. Dark green: Failed relationships, e.g.-drug failed to treat a disease. For fully interactive figure, visit: https://github.com/gtsueng/M2C_rel_nb/tree/master/results/no_relation_broad_categories.html

**Supplemental Figure 2-.**
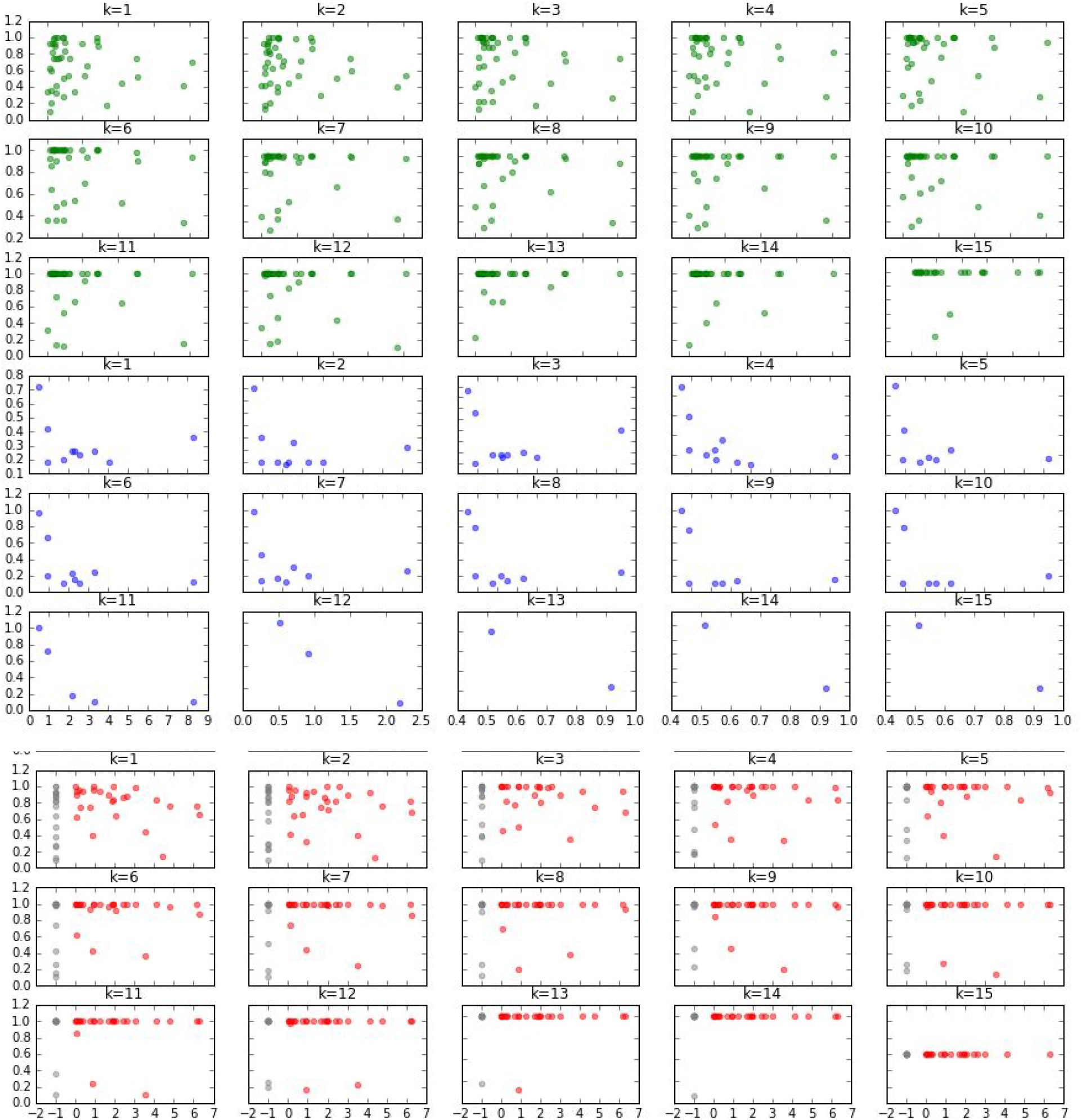
The effect of concept distance on accuracy at different levels of K. Concept Distance vs Accuracy of Related (green) or Unrelated Concept (blue) Pairs or ‘Broken Concepts’ (red). In grey are concepts with no identifiers for which concept distance could not be calculated. Accuracy appears to be more greatly affected by voter threshold (k) than concept distance.

